# Single cell long read genotyping of transcripts reveals discrete mechanisms of clonal evolution in post-MPN AML

**DOI:** 10.1101/2025.08.18.670417

**Authors:** Julian Grabek, Jasmin Straube, Leanne Cooper, Rohit Haldar, Ranran Zhang, Inken Dulige, Matthew Barker, Will Gatehouse, Helen Christensen, Gerlinda Amor, Victoria Y. Ling, Caroline McNamara, David M. Ross, Andrew Perkins, Megan J. Bywater, Steven W. Lane

## Abstract

Myeloproliferative neoplasms (MPNs) are caused by acquired mutations in hematopoietic stem and progenitor cells (HSPCs). The acquisition of additional mutations like *TP53* and the overall mutational burden influence a patient’s risk of disease progression toward lethal post-MPN acute myeloid leukemia (AML).

Recent technological advancements in linking single-cell gene expression with genotype have improved our understanding of tumor heterogeneity. However, current methodologies have limitations in simultaneously genotyping low-expression genes (such as *JAK2*) alongside other pathogenic loci.

To address this, we developed a novel <u>lo</u>ng read genotyping pipeline of cDNA <u>tr</u>anscripts called LOTR-Seq, which can genotype the full length of expressed transcripts of 30 genes at once. Using LOTR-Seq, we genotyped HSPCs at the *JAK2*V617 locus in 9,075 single cells from eight patients with chronic phase MPN (CP-MPN) and in 5,016 cells from four patients with post-MPN AML. We then linked the mutations to the single cell transcriptome of 29,712 *JAK2*V617F-driven CP-MPN cells and 16,895 post-MPN AML cells.

In our analysis of post-MPN AMLs, we identified nine mutated loci across six genes (*JAK2, IDH1/2, TP53, SRSF2, U2AF1*) and linked these mutations to specific transcriptional phenotypes. Overall, LOTR-Seq provides novel insights into the evolution of post-MPN AML.

## Introduction

Myeloproliferative Neoplasms (MPN) are clonal blood cancers characterized by the overproduction of mature, functional myeloid elements. MPNs are caused by driver mutations within the hematopoietic stem and progenitor (HSPC) compartment^1^, with the most common being *JAK2*V617F^2^. *JAK2*V617F gives rise to a clonal stem cell population^3^ and results in pathology mediated by mature myeloid cells. Both healthy and MPN HSPCs exhibit both transcriptional evidence of myeloid and lymphoid lineage priming^4^ and self-renewal capacity^5^. Exactly how this clonal outgrowth affects the transcriptional diversity of the HSPC compartment is unclear.

In contrast to MPN, acute myeloid leukemias (AML) are characterized by the overproduction of immature myeloid cells. Genetic complexity is a common feature of *JAK2*V617F post-MPN AML, with patients having multiple additional mutations identified at AML transformation^6^. Disease progression from chronic phase MPN (CP-MPN) to AML is variable and presumed to be dictated by the acquisition of mutations in HSPCs in addition to the MPN driver and subsequent clonal expansion^7,8^.

Single cell technologies have provided unique insights behind this evolution, however current methods are constrained by the cell number, and the number of genetic loci that can be concurrently analyzed (Supplemental Figure 1)^7–10^. We have developed an innovative pipeline called LOTR-Seq, which allows for concurrent single-cell transcription plus genotyping of a comprehensive panel of full-length transcripts from 30 genes that are recurrently mutated in post-MPN AML. This full-length approach facilitates the detection of rare variants, the phasing of mutations, and the identification of different isoforms.

Here we used LOTR-Seq to identify clonal evolution during transformation from MPN to AML and to determine the impact of these additional mutations on *JAK2*-mutant HSPCs in driving disease progression.

## Materials and methods

### Ethics approval and Primary Human Sample processing

Patient bone marrow (BM) samples were obtained from the Royal Brisbane Women’s Hospital and from the South Australian Cancer Research Biobank (SACRB), and analyzed with approvals from QIMR Berghofer HREC (p1382). Patients provided written informed consent for the collection, storage, and use of their samples in research. Detailed patient information can be found in Supplemental Table 1. Samples were stained with CD3-PE/Cy7, CD34-PE, CD14-FITC and CD15-APC (Supplemental Table 2). CD34+ HSPCs were FACS sorted, and single cell 3’ RNA sequencing was performed with the 10x Chromium v3.1 on the Illumina NextSeq 500/550 Platform. CD34+ BM bulk genomic DNA capture panel sequencing and mutation calling were performed as described in Magor et al, 2016^11^ on 7 of the 12 samples.

### Long read sequencing of transcripts: LOTR-Seq

We first designed a capture panel on the Roche HyperDesign platform targeting all exonic regions of 30 recurrently mutated MPN/AML genes: *ASXL1*, *BCOR*, *CALR*, *CBL*, *CSF3R*, *DNMT3A*, *EZH2*, *FLT3*, *GATA2*, *IDH1*, *IDH2*, *JAK2*, *KIT*, *KRAS*, *MPL*, *MYC*, *NPM1*, *NRAS*, *PPM1D*, *PTPN11*, *RUNX1*, *SF3B1*, *SH2B3*, *SRSF2*, *STAG2*, *TET2*, *TP53*, *TERT*, *U2AF1,* and *ZRSR2*. 10x barcoded cDNA underwent pre-capture PCR and read extension using custom primers (Supplemental Table 3)^12^. Amplified and extended cDNA was used to capture genes of interest using the default KAPA HyperCap Workflow v3, followed by post-capture PCR amplification. Further detailed information can be found in the Supplemental methods. Oxford Nanopore Technology (ONT) ligation kits, LSK110 or LSK114, were used in accordance with the manufacturer’s protocols, with modifications to the incubation time from 10 minutes to 20 minutes^13^ to improve adapter ligation and ONT Flow Cell loading amount (50-100 fmol; Supplemental Table 4).

### Single cell short read sequencing processing

MPN and post-MPN AML sequenced reads were adapter trimmed and processed through cellranger (v6.0.1, or MPN10 v3.0.2) with GRCh38 genome build to obtain a counts per single cell and gene matrix. 10x single cell RNA-seq data from healthy human CD34+ BM HSPCs from Ainciburu et al. 2023^14^ were downloaded from GEO (Supplemental Table 5). Healthy, CP-MPN, and post-MPN AML samples were preprocessed in R (v3.6.3) following the Seurat v4.3 pipline^15^. Clusters were annotated according to key lineage marker gene expression. For analyzing lineage skewing, clusters were broadly grouped by adding up clusters into HSC/LMPP (HSC+LMPP), Mega/Ery-Primed (MEP+MkP+e-Ery+l-Ery+Ery), Gran/Mono-Primed (GMP+Mono+Baso+DC), and Lymphoid-Primed (CLP+B-primed). Pseudo time prediction and cell fate scoring were performed using Palantir (v1.3.0)^16^ within Python (v3.11.4). Differential gene expression and gene set enrichment analysis are detailed in Supplemental methods.

### ONT sequencing, read processing and mutational profiling analysis

Base calling and chimeric read separation are detailed in Supplemental methods. Reads were assigned a 10x barcode using FLAMES^17^, allowing for two mismatches. Fastqs were split into individual barcode fastqs using zgrep to identify FLAMES-assigned barcodes in the fastq header line. Deconvoluted reads were mapped against the GRCh38 genome build using minimap2 (v2.27)^18^ with option ‘-L -ax splicè. Reads filtering and mutation calling are detailed in the Supplemental methods. Expressed variant allele frequency (eVAF) was calculated in R as the ratio of the number of reads with bases matching the mutant allele to the total number of reads at the locus. Finally, eVAFs were assigned to a single cell transcriptome by matching barcodes recovered by long read and short read sequencing.

### Post-MPN AML differentiation stage assessment

ScRNASeq raw counts from 11 post-MPN AML *JAK2*V617F*/TP53* bi-allelic loss^8^ and our post-MPN AML samples were projected on the Bone marrow map as described in https://github.com/andygxzeng/BoneMarrowMap^21^ (accessed 27/04/2025). Raw counts of bulk RNA-Seq data from 45 CP MPN, 34 post-MPN AML, and 11 healthy controls were downloaded from GEO with GSE283710^20^. Single cell RNA-Seq (scRNASeq) data of 11 *JAK2*V617F*/TP53 bi-allelic loss* post-MPN AML and 5 healthy controls (GSE226340)^8^ were extracted and raw counts pseudo-bulked. Data were normalized (edgeR v4.2.2) and log2 transformed before differentiation stage scoring was performed as described^21^. AML differentiation scores from bulked and pseudo-bulked RNASeq data were centered, and patient-specific scores Euclidean distance underwent hierarchical clustering (hclust; stats R base package) with default parameters. Data were visualized with heatmap3.R https://github.com/obigriffith/biostar-tutorials/blob/master/Heatmaps/heatmap.3.R.

## Results

### HSPC heterogeneity is preserved in chronic phase MPN but with lineage skewing

To determine whether the expression of *JAK2*V617F alters the transcriptional heterogeneity of the CD34+ HSPC compartment, we compared the single cell transcriptional landscape of CD34+ HSPC-enriched populations (Figure 1A) from normal human bone marrow samples^14^ (Supplemental Figure 2A; n=6) to that of CD34+ HSPCs from eight patients with *JAK2*V617F-driven CP-MPN (Figure 1B). Transcriptional heterogeneity was largely preserved in CP-MPN CD34+ cell populations, comprising HSC/LMPPs, megakaryocyte-erythroid-, lymphoid- or granulocyte-monocyte-primed populations, and consistent with that seen in the healthy CD34+ populations (Figure 1B-C; Supplemental Figure 2B). However, CP-MPN CD34+ populations demonstrated a higher percentage of cells exhibiting megakaryocyte/erythroid-primed gene expression profiles (Figure 1D-E: Mega/Ery-Primed mean±SD: 22.5±6.9% healthy vs 34.1±10.3% CP-MPN, p=0.028; Supplemental Figure 2C).

**Figure 1.**
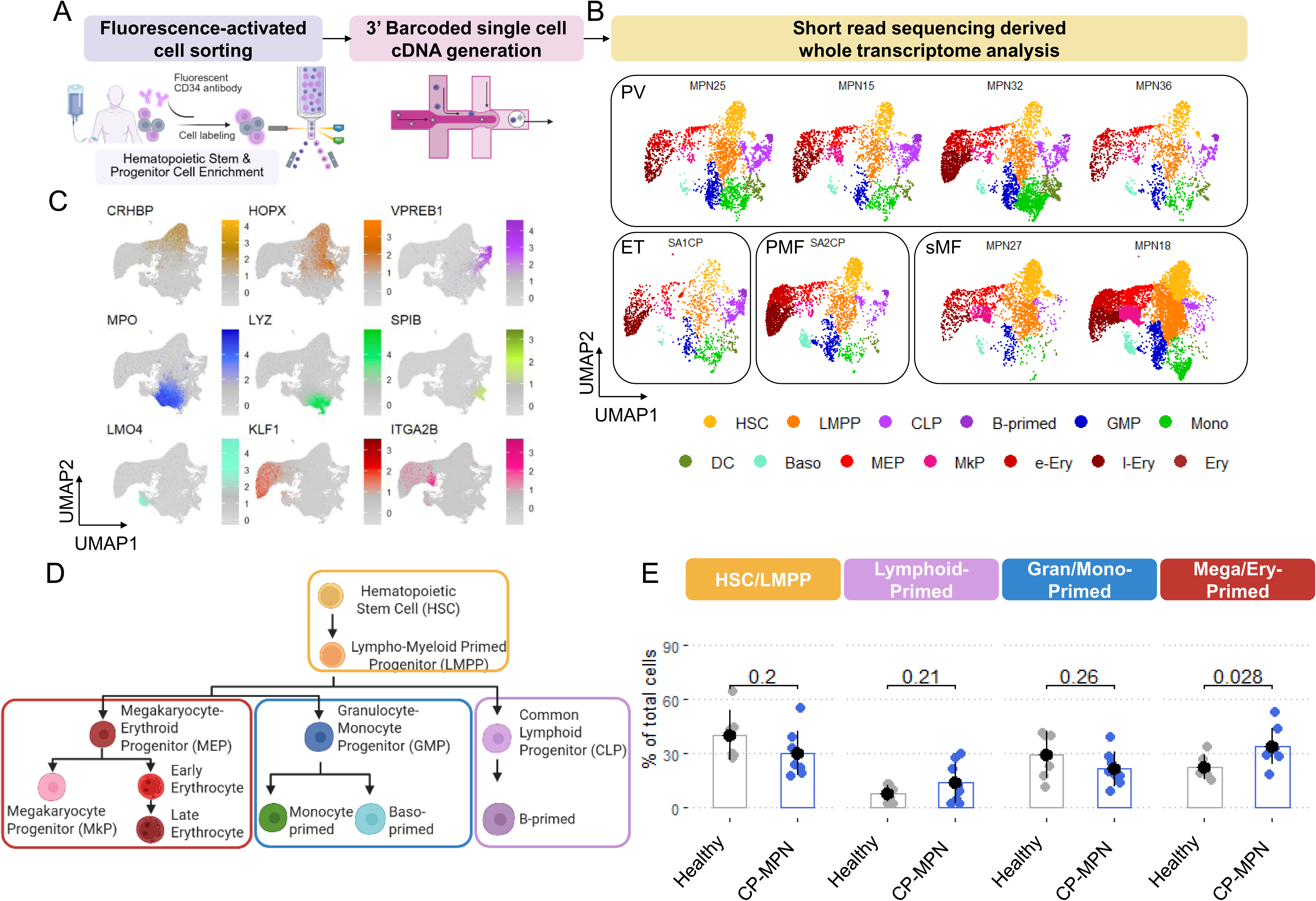
**Single cell RNA sequencing identifies expanded megakaryocyte-erythroid primed cells in chronic phase MPN.** (A) Schematic of MPN bone marrow hematopoietic stem and progenitor cell (HSPC) enrichment followed by 10x Chromium single cell separation and 3’ cDNA barcoding. (B) Chronic phase MPN UMAPs (n=4, polycythemia vera, PV; n=1, essential thrombocythemia, ET; n=1 primary myelofibrosis, PMF; n=2 secondary myelofibrosis, sMF) of short read sequencing derived whole transcriptomics of HSPCs. Clusters annotated by gene expression associated with HSC, hematopoietic stem cells; LMPP, lympho-myeloid multipotent progenitor; CLP, common lymphoid progenitors; GMP, granulocyte-monocyte progenitors; DC; Dendritic Cell progenitors; Baso, eosinophil/basophil/mast cells progenitors; MEP, megakaryocyte-erythroid progenitor; MkP, megakaryocyte progenitor, e-Ery; early erythroid and l-Ery, late erythroid priming. (C) Pooled healthy sample UMAPs (n=6) showing the expression of key lineage-defining transcriptional markers including HSC-CRHBP, LMPP-HOPX, B-primed-VPREB1, myeloid-MPO, monocyte-LYZ, DC-SPIB, basophil-LMO4, erythroid-KLF1 and megakaryocyte-ITGA2B. (D) Schematic of summarized clusters analysed in (E). (E) Bargraph with mean (black dot) and standard deviation (black line) comparing the percentage (%) of clustered cells in healthy (n=6) vs chronic phase (CP) MPN (n=8). Each grey or blue dot represents data from an individual sample. P-value derived from pairwise, two-sided Welch t-test. (A, D) Created in BioRender. Lane, S. (2025) (A) https://BioRender.com/vygkh71, (D) https://BioRender.com/9t72apn).

These findings demonstrate that, consistent with healthy CD34+ HSPCs, transcriptional lineage-priming is present in primitive hematopoietic cell populations in CP-MPN; however, CP-MPN exhibits an HSPC compartment with a higher abundance of cells with a megakaryocyte/erythroid transcriptional phenotype, consistent with the pathology of CP-MPN, characterized by the overproduction of mature megakaryocytes and erythrocytes.

### MEP lineage priming in HSPCs is linked to the expansion of *JAK2*V617F mutant cells

We were interested to determine how clonal heterogeneity within CP-MPN altered this lineage bias within the CD34+ HSPC population. To identify *JAK2*V617F-mutated cells within CP-MPN CD34+ HSPCs, we developed a technique for long read genotyping of transcripts (LOTR-Seq). LOTR-Seq uses the single cell barcoded cDNA derived by the 10x Chromium platform to capture the cDNA of target genes, like *JAK2*, followed by ONT long read sequencing (Figure 2A) to correlate expressed mutational information to the transcriptome.

**Figure 2.**
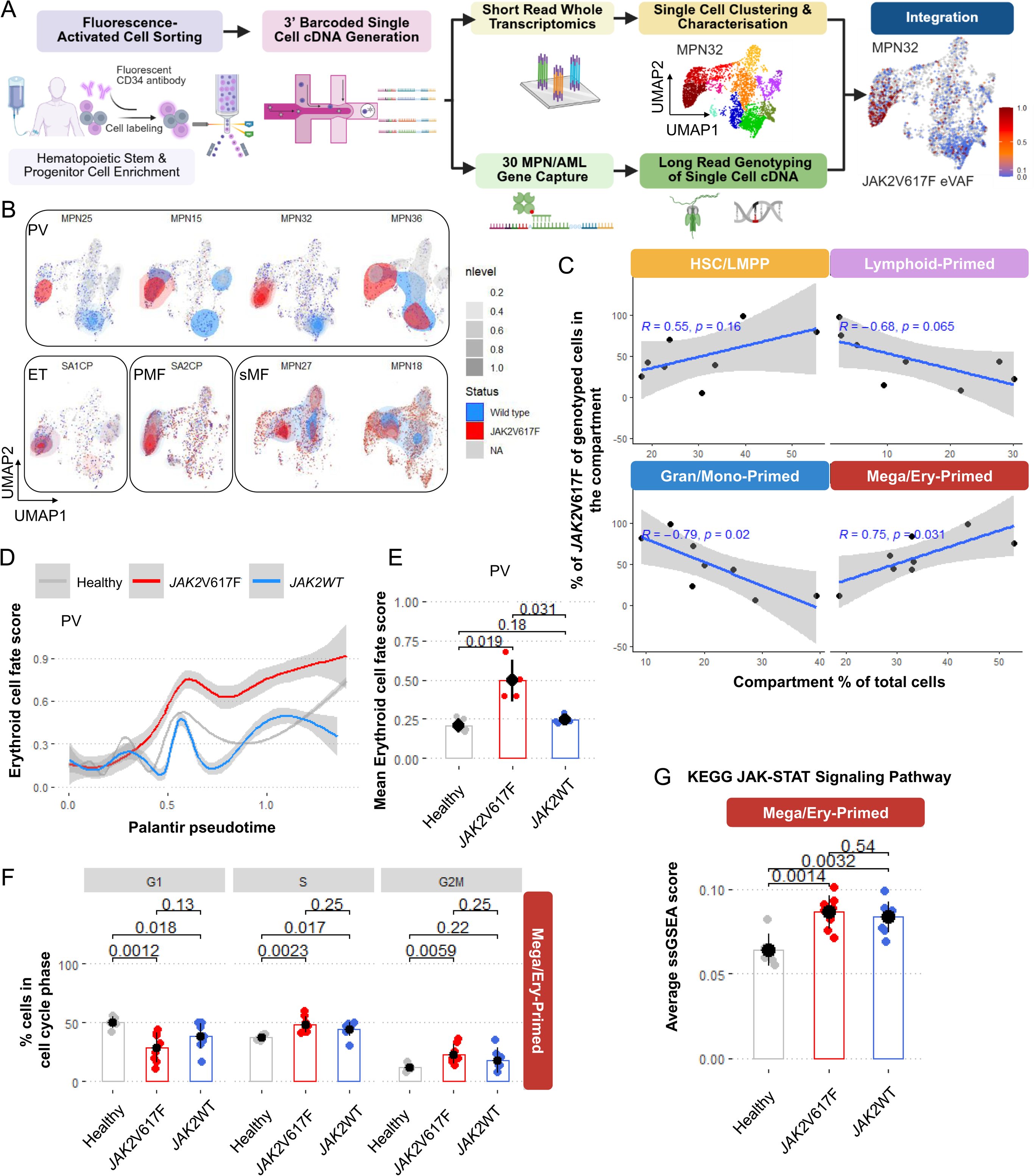
**Target enrichment and long read sequencing demonstrates the impact of mutational status on the gene transcription in chronic phase MPN.** (A) Schematic showing the long read genotyping of transcripts (LOTR-Seq) pipeline that enriches single cell cDNA of 30 MPN/AML associated genes followed by long read sequencing and genotyping of transcripts, with an example of one MPN patient’s UMAP colored by *JAK2*V617F expressed variant allele frequency (eVAF). (A) Created in BioRender. Lane, S. (2025) https://BioRender.com/vygkh71. (B) UMAPs of the CD34+ HSPC single cells (light grey) of n=8 MPNs with the density distribution of *JAK2*V617F mutation (red), *JAK2*-wild type (blue) and non-genotyped cells (NA; grey). (C) Scatterplots of HSC/LMPP and key lineage-primed compartment percentage (%; x-axis) and the % of *JAK2*V617F genotyped cells (y-axis) within the compartment (n=8 chronic phase MPN). Pearson’s correlation coefficient R and test of association p-value. The blue line represents the linear regression line with a 95%-confidence interval (grey band). (D) Smoothed regression lines of the Palantir predicted erythroid cell fate score (y-axis) in dependency of the pseudotime (x-axis) for n=5 healthy, n=4 Polycythemia vera patients (PV) *JAK2*V617F and *JAK2* wild type cells. (E) Bargraph of the mean erythroid cell fate for each sample comparing n=5 healthy to n=4 PV *JAK2*V617F and *JAK2*-wild type (*JAK2*WT) summarized in (D). Black dot and line represent the group mean and standard deviation, respectively. P-values derived from pairwise, two-sided Welch t-tests. (F) Bargraph of the percentage (%) of cell cycle phase (G1, S, G2M) for the Mega/Ery-Primed compartment comparing n=6 healthy to n=8 CP-MPN *JAK2*V617F and *JAK2* wild type (*JAK2*WT). Black dot and line represent the group mean and standard deviation, respectively. P-values derived from pairwise, two-sided Welch t-tests. (G) Bargraph of the KEGG JAK-STAT signaling pathway average single cell gene set enrichment score comparing n=6 healthy to n=8 MPN *JAK2*V617F and *JAK2* wild type (*JAK2*WT). Black dot and line represent the group mean and standard deviation, respectively. P-values derived from pairwise, two-sided Welch t-tests.

When LOTR-Seq was performed on the CP-MPN CD34+ HSPCs, we were able to demonstrate that *JAK2*V617F pseudo-bulked eVAF strongly correlated with both genomic VAF (r=0.9; p=0.005; Supplemental Figure 3A) and short read-derived eVAF (r=0.96; p<000.1; Supplemental Figure 3B) of the CD34+ compartment, validating the reproducibility of mutation calling from long read sequencing. LOTR-Seq enabled us to genotype up to 51% of cells at the *JAK2*V617 locus, improving upon previous techniques (Supplemental Figure 1)^10,22^. Notably, the percentage of genotyped cells at the *JAK2*V617 locus correlated with sequencing depth (r=0.8, p=0.0006; Supplemental Figure 3C).

We then assessed the contribution of *JAK2*V617F-mutated cells to the CD34+ HSPC compartment using the density distribution of *JAK2*V617F-mutated and *JAK2*-wild type cells within the clusters identified by uniform manifold approximation and projection (UMAP) (Figure 2B). The percentage of *JAK2*V617F mutated HSPCs within the Mega/Ery-primed CD34+ compartment correlated with the relative size of the Mega/Ery-primed compartment (R=0.75; p=0.031; Figure 2C) and inversely correlated with that of the Gran/Mono-primed compartment (R=-0.79; p=0.02). PV patients displayed higher *JAK2*V617F clone percentages in the Mega/Ery-Primed compartments^10^ (Supplemental Figure 3D), conversely, *JAK2*V617F lineage quantification was stably distributed in patients with ET or MF.

These findings suggest that *JAK2*V617F either skews lineage bias towards the Mega/Ery fate or confers a proliferative advantage exclusively to Mega/Ery-primed HSPCs. To resolve this, we used Palantir analysis of differentiation in pseudotime to examine the effect of *JAK2*V617F on HSPC cell fate potential^16^. Here, HSPCs of PV patients containing the *JAK2*V617F mutation show a higher probability of converging towards an erythroid cell fate, evidenced by a higher average erythroid cell fate score compared to both healthy CD34+ and *JAK2*-wild type cells (Figure 2D-E, Supplemental Figure 3E), while sMF patients exhibit a higher probability of converging towards a megakaryocytic progenitor fate (Supplemental Figure 3F-G). This supports the finding that *JAK2*V617F favors erythroid/megakaryocyte differentiation. Furthermore, cell cycle analysis revealed that *JAK2*V617F mutant cells contained a greater proportion of cells in G2M phase (Figure 2F, Supplemental Figure 4A).

We next sought to determine the impact of the *JAK2*V617F mutation on canonical signaling pathway activation. Compared to healthy CD34+, CP-MPN CD34+ demonstrated increased JAK-STAT signaling across all compartments (Figure 2G, Supplemental Figure 4B) in both *JAK2*V617F, *JAK2*-wild type cells, suggesting that the activation of JAK-STAT signaling within HSPC is also regulated by cell-extrinsic factors, such as increased activation of cytokine pathways^23^. The impact of this *JAK2*V617F mutation appears most pronounced in the Mega/Ery-Primed compartment with Myc targets and the heme metabolism pathway activated in CP-MPN. Interestingly, inflammatory pathways are suppressed in PV but activated in MF patients, particularly at the HSC/LMPP level (Supplemental Figure 4C).

These data validate the LOTR-Seq pipeline to identify *JAK2*V617F mutant vs *JAK2*-wild type HSPCs within individual CP-MPN samples. *JAK2*V617F mutated cells demonstrate increased proliferation and preferential expansion towards the erythroid and megakaryocyte lineages and context-specific transcriptional effects of *JAK2*V617F by lineage-primed compartment and MPN subtype, reflecting the heterogeneity observed in the clinical presentation of MPNs.

### Transformation to post-MPN AML is characterized by the loss of stem cell transcriptional heterogeneity and is driven by genetic complexity

We next sought to determine whether the LOTR-Seq pipeline could be used to investigate the relationship between genetic and transcriptional heterogeneity within the HSPC compartment during progression from CP-MPN to post-MPN AML. In dramatic contrast to the transcriptional heterogeneity observed in CP-MPN, the CD34+ HSPC compartment in post-MPN AML is largely dominated by a single transcriptional profile (Figure 3A-B). For example, MPN10 AML exhibits a dominant HSC/LMPP-like transcriptional signature, representing 79.4% of all cells (Figure 3A), while MPN20 AML shows a dominant Mega/Ery primed-like transcriptional signature (84.8%; Figure 3B).

**Figure 3.**
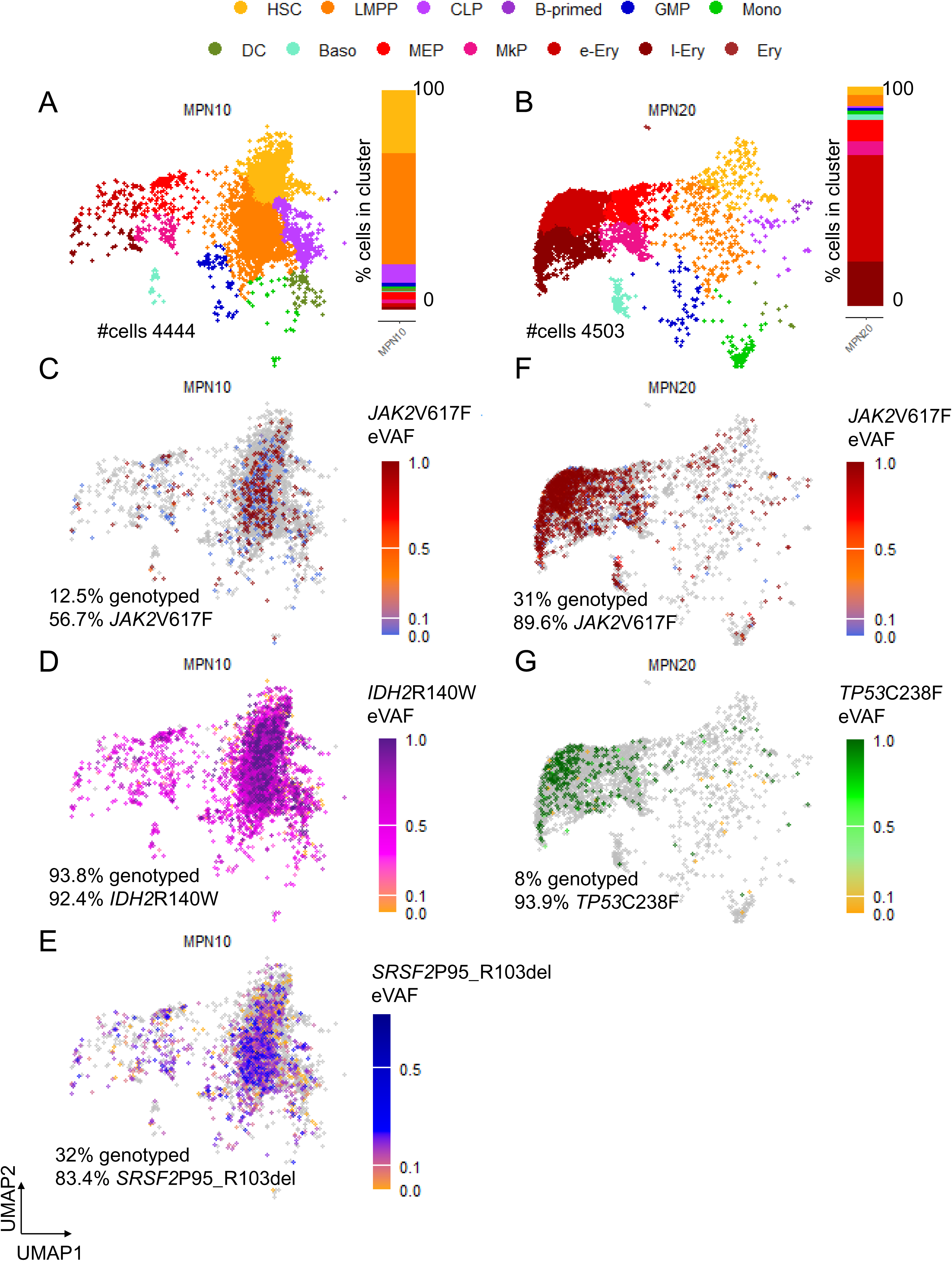
**Loss of heterogeneity occurs in leukemic transformation and is driven by additional mutations in the *JAK2*V617F mutant clone.** (A-B) UMAPs of post-MPN AML samples (A) MPN10 and (B) MPN20 showing transcriptionally defined cells clusters and a bargraph depicting the relative percentage of cells per cluster, (C-E) UMAPs of post-MPN AML sample MPN10 showing expressed variant allele frequencies (eVAF) of mutations (C) *JAK2*V617F (D) *IDH2*R140W and (E) *SRSF2*P95_R103del. (F-G) UMAPs of post-MPN AML sample MPN20 showing expressed variant allele frequencies (eVAF) of mutations (F) *JAK2*V617F and (G) *TP53*C238F. Grey colored dots indicate non-genotyped cells.

A limitation of existing protocols integrating single cell RNA-Seq and genotyping is the inability to simultaneously genotype across numerous loci with sufficient coverage and representation to draw inferences on cells with multiple mutations within the CD34+HSPC compartment (Supplemental Figure 1)^22^. The LOTR-Seq pipeline overcomes this limitation by using enrichment for a panel of 30 genes, enabling the genotyping of expressed genes across a range from 0% to 100% (Supplemental Figure 5A). We therefore sought to apply this technology to determine the relationship between the genomic architecture of post-MPN AML and the transcriptional consequences linked with disease progression.

MPN10 contained a dominant, HSC/LMPP-like expanded population in post-MPN AML. LOTR-Seq identified the MPN-driver *JAK2*V617F mutation together with an *IDH2*R140W mutation and *SRSF2*P95_R103 in-frame deletion (Figure 3A, C-E, Supplemental Figure 5B) in 56.7%, 92.4 %, and 83.4% of genotyped cells, respectively, and expressed at high VAF in the HSC/LMPP-like CD34+ cells. Additionally, in *JAK2*V617F, *IDH2*R140W and *SRSF2*P95R co-mutated cells we observed the predominant inclusion of a poison exon cassette^24^, in comparison to cells from a post-MPN AML containing *JAK2*V617F and *IDH1*R132S without a mutation in *SRSF2* (Supplemental Figure 5C).

In a second post-MPN AML, MPN20, we identified a dominant Mega/Ery primed-like HSPC population containing *JAK2*V617F (89.6% of genotyped cells) with biallelic *TP53*C238F mutation (93.9% of genotyped cells) (Figure 3B, F-G, Supplemental Figure 5D). This identification of a Mega/Ery-signature in *JAK2*V617F multi-hit-*TP53* post-MPN AML is consistent with other published human data^8^ and aligns with data from murine post-MPN AML with this same combination of mutations^25,26^.

Altogether, these data validate the use of LOTR-Seq in clonal analysis of human post-MPN AML and demonstrate that post-MPN AMLs exhibit loss of HSPC heterogeneity with the expansion of a dominant transcriptional cluster that appears to be linked to the identity of co-existing secondary mutations.

**Post-MPN AML demonstrates lineage restricted, recurrent differentiation priming** Recent findings suggest that molecular subtypes of de novo AML are defined by unique transcriptional profiles that reflect lineage bias^21,27,28^. Our initial analysis of the HSPC compartment in post-MPN AML suggests lineage priming analogous to that seen in de novo AML^21,28^. To determine if these findings can be applied more generally to post-MPN AML, we analyzed bulk RNA-Seq of 34 CD34+ post-MPN AML samples^20^ together with single cell RNA-Seq CD34+ HSPCs of a further 11 post-MPN AML cases^8^. Here, we adopted methods developed by Zeng et al 2025^21^ to assess differentiation stages in post-MPN AML. Both methods of projecting AML on a detailed, healthy bone marrow hierarchical hematopoiesis map^21^ (Figure 4A-B) and AML differentiation stage scoring (Figure 4C) suggest that even genetically homogenous subgroups, such as *JAK2V617F* with multi-hit *TP53*, can exhibit expansion in different lineage-primed compartments. Compared to adult de novo AML, as reported by Zeng et al 2025^21^ (n=28 of 139), post-MPN AML exhibited a significantly higher overrepresentation of HSC/LMPP states (n=23 of 45, Chi-Square p=0.0001). In the 45 post-MPN AML samples, 51% showed an immature differentiation stage (HSC/LMPP/MEP), with HSC/LMPP/early lymphoid stage or MEP/MKP/erythroid stage observed in 31% and 18%, respectively. Strikingly, there was an absence of a promonocytic/monocytic biased subgroup in *JAK2*V617F mutated post-MPN AML, a dominant subgroup in de novo AML^21^ (Figure 4C). These data demonstrate that post-MPN AML is not only genetically distinct from de novo AML, but also phenotypically and transcriptionally distinct with a more restricted differentiation repertoire.

**Figure 4.**
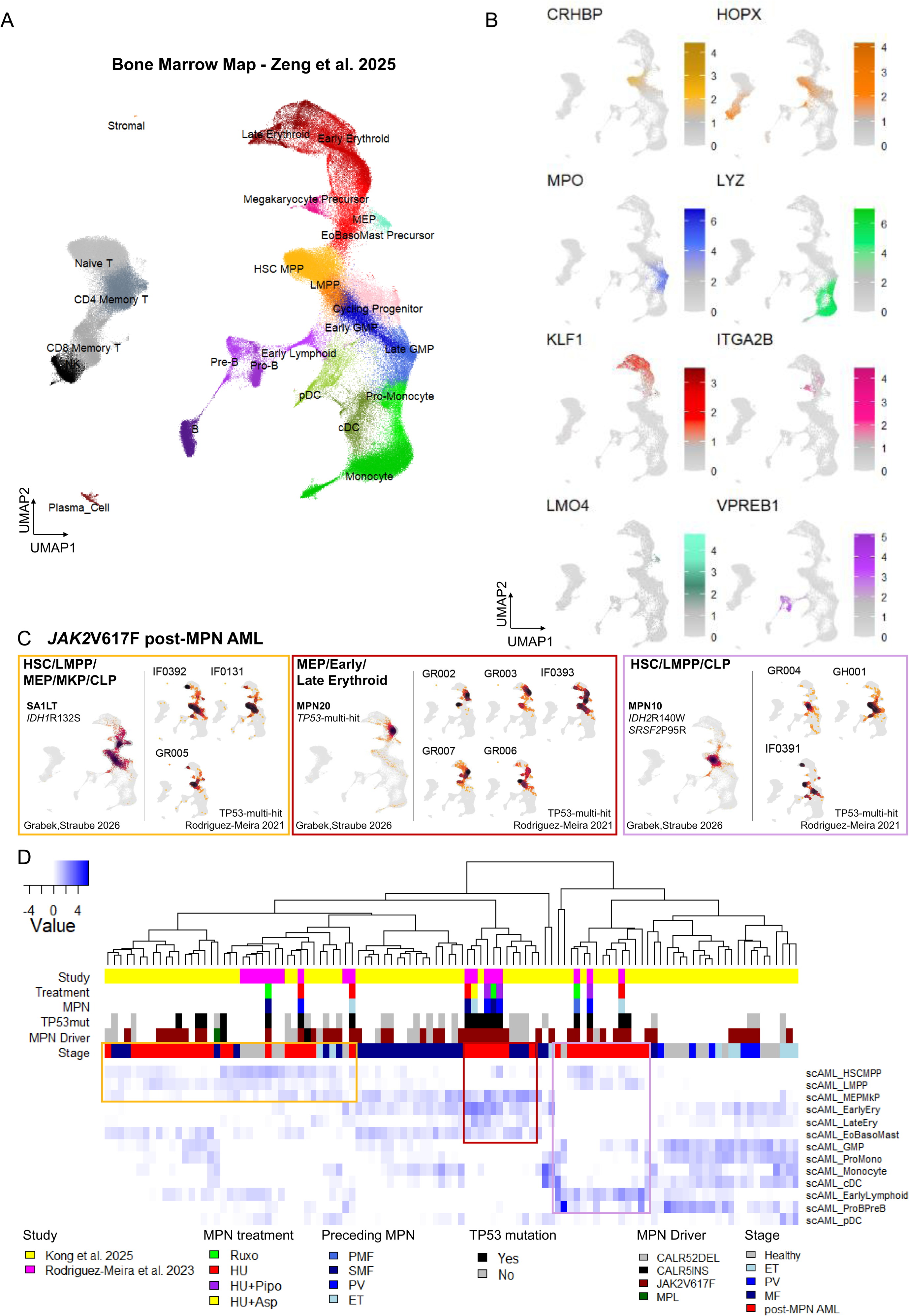
**Post-MPN AML exhibit lineage-biased hematopoiesis** (A) Bone marrow hematopoietic stem cell atlas UMAP with cell type annotations as generated by Zeng et al. 2025^21^. (B) UMAPs highlighting key lineage gene expression markers HSC-*CRHBP*, LMPP-*HOPX*, erythroid-*KLF1*, megakaryocyte-*ITGA2B*, Baso-*LMO4*, and B-primed-*VPREB1*. (C) Projection of our generated n=3 *JAK2*V617F post-MPN AML single cell samples (SA2LT, MPN20, MPN10)and n=11 *JAK2*V617F TP53-multi-hit post-MPN AMLs from Rodriguez-Meira et al. 2023^8^ on to the healthy bone marrow atlas UMAP. (D) Hierarchical clustering of lineage-defining gene expression scores (y-axis) of n= 11 *JAK2*V617F post-MPN AML and n=6 healthy samples from Rodriguez-Meira et al. 2023^8^, n=45 chronic phase MPN, n=34 post-MPN AML and n=11 healthy from Kong et al 2025^20^ (x-axis).

### High-risk mutations in post-MPN AML are already present in chronic phase disease

Post-MPN AML is intrinsically chemotherapy resistant, and novel strategies are urgently required to prevent the progression of high-risk CP-MPN to AML. We therefore sought to determine whether LOTR-Seq could identify pre-leukemic high-risk populations within CD34+ cells from patients with CP-MPN. To do this, we analyzed a patient with matched CP-MPN and post-MPN AML samples (Figure 5). The *JAK2*V617F mutation was stable between CP-MPN and post-MPN AML (Figure 5B, E), however deep sequencing analysis of the CP-MPN CD34+ cells identified a rare subpopulation of co-mutated *JAK2*V617F and *IDH1*R132S (8.2%) in CP-MPN, which expanded to 64.5% in post-MPN AML (Figure 5C, F). In contrast to what was observed in the analysis of the previous post-MPN AML samples (Figure 4), the expansion of the double-mutated population resulted in the relative expansion of the HSC/LMPP clusters with partially maintained representation of the more committed lineage-primed cells.

**Figure 5.**
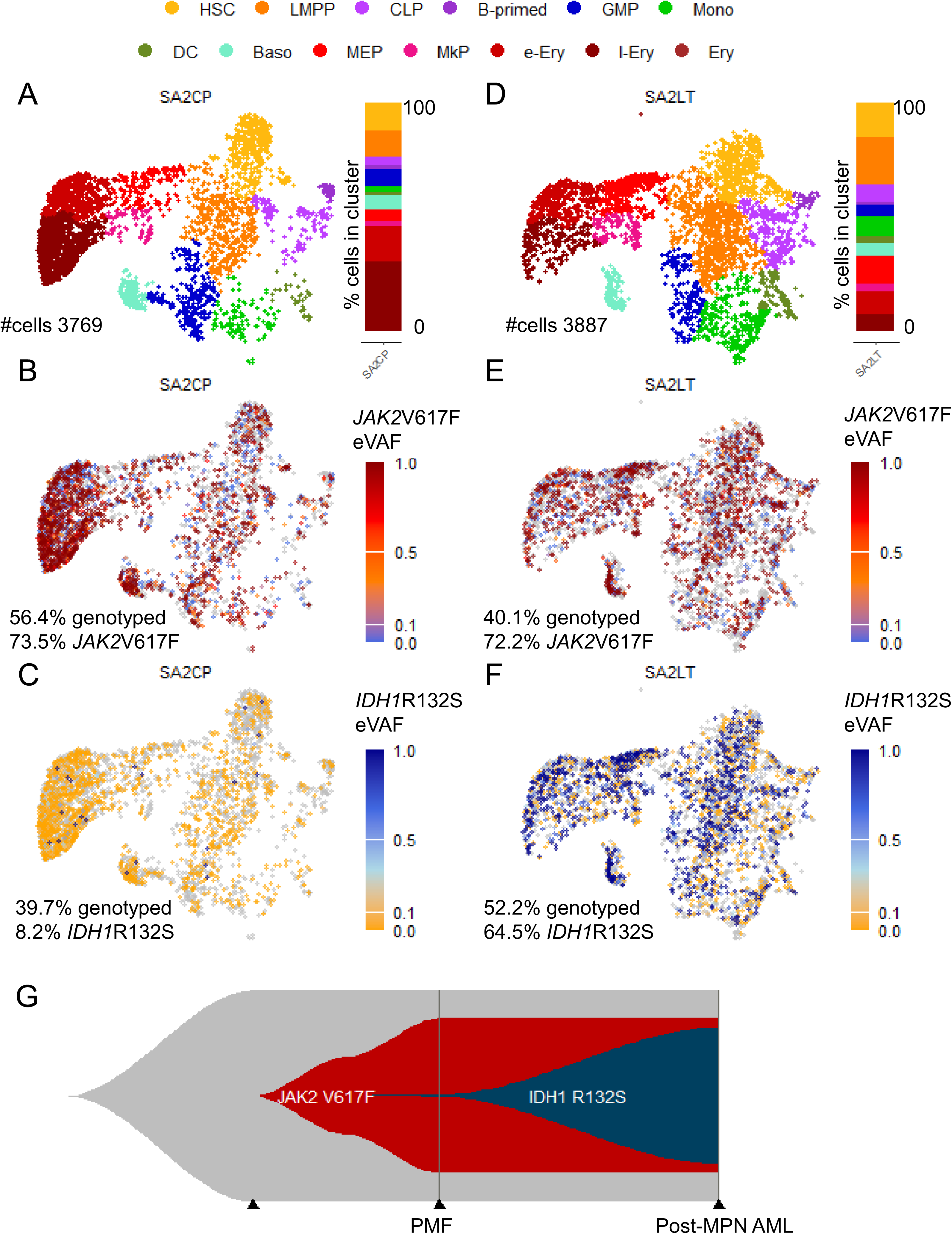
**Leukemic transformation from chronic phase MPN showing a linear pattern of clonal expansion of a high risk *JAK2*V617F and *IDH1*R132S mutant clone.** UMAPs of the same patient at diagnosis of (A-C) chronic phase MPN (SA2CP) and (D-F) post-MPN AML (SA2LT), showing (A,D) transcriptionally defined clusters with bargraphs showing the percentage of cells in the transcriptionally defined clusters and (B-C,E-F) expressed variant allele frequencies (eVAF) of mutations (B,E) *JAK2*V617F and (C,F) *IDH1*R132S. Grey colored dots indicate non-genotyped cells. (G) Fish plot illustrating the linear clonal evolution of the *JAK2*V617F and *IDH1*R132S mutant clone.

The co-occurrence of *JAK2*V617F and *IDH1*R132S clones in chronic phase, with expansion of the double-mutated clone during AML progression, suggests that this leukemia was driven by the linear clonal evolution of the double-mutant cells (Figure 5G). Thus, the LOTR-Seq protocol can identify genetic complexity at the single cell level, enabling the identification of pre-leukemic clones that later give rise to post-MPN AML.

### Additional mutations in *JAK2*V617F cells exhibit mutation and context specific gene expression signatures

To determine how the presence of additional mutations can impact MPN stem cells and if this changes with leukemic transformation, we first compared *IDH1*R132-mutant cells (n=122) with *JAK2*V617F/*IDH1*WT cells (n=1562), in the matched chronic phase sample, for differential gene expression and gene set enrichment across the major transcriptional compartments in CD34⁺ bone marrow (Figure 6A-C, Supplemental Table 9-10). *IDH1*R132S cells within the Mega/Ery-Primed compartment displayed heightened activation of inflammatory pathways, notably TNFα and IL2–STAT5A, alongside an increase in *STAT5A* gene expression, known to have potential roles in MPN pathophysiology^20,29^. Concurrently, we observed an upregulation of hypoxia-related genes (e.g., *BCL2*) and a downregulation of oxidative phosphorylation (OXPHOS)-related genes, suggesting a metabolic shift in the *IDH1*-mutant cellular environment.

**Figure 6.**
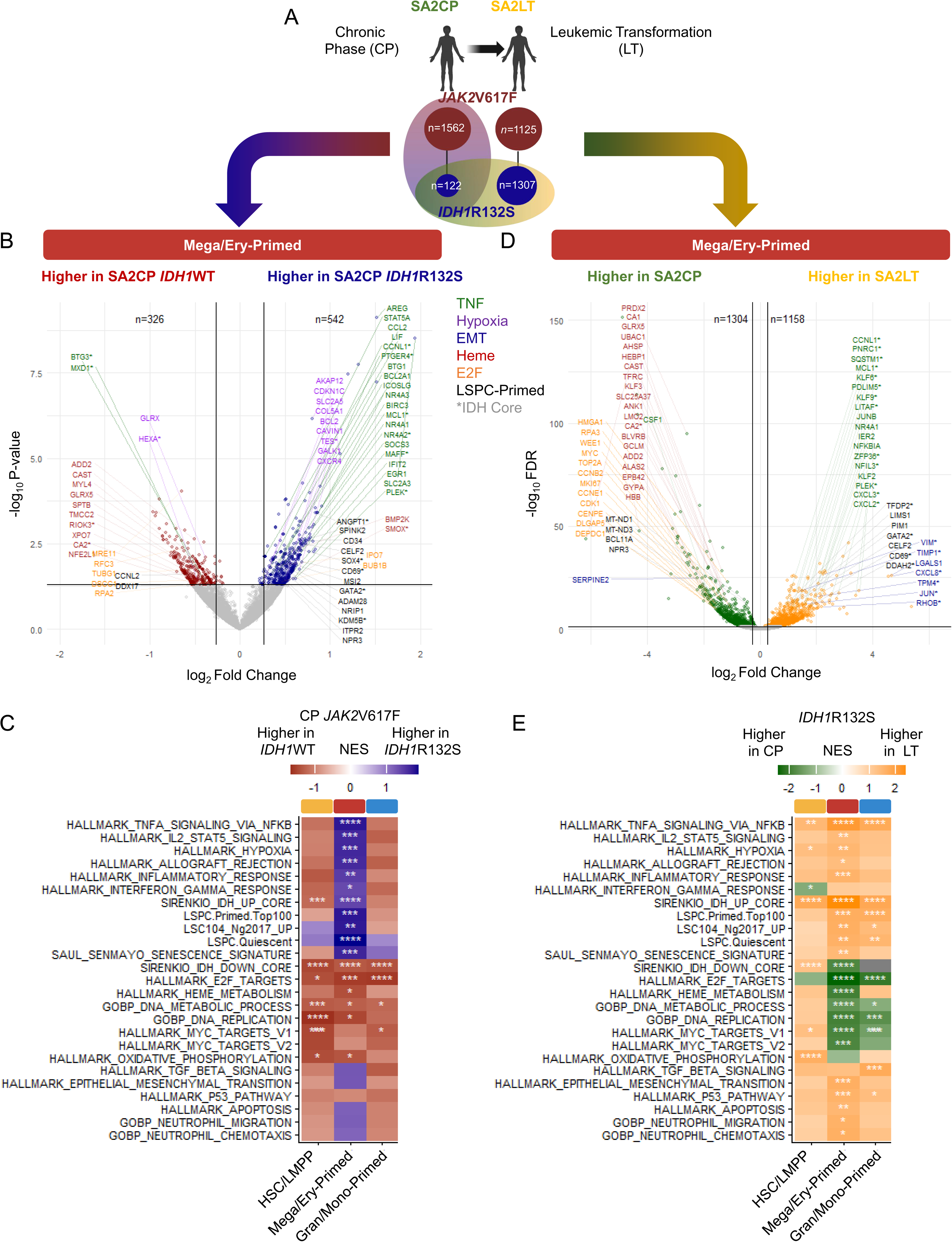
**IDH1 mutations in *JAK2*V617F cells exhibit mutation and context specific gene expression signatures** (A) Schema of analysis performed in B-E, Created in BioRender. Lane, S. (2026) https://BioRender.com/1kuev8s (B) Volcanoplot displaying the log_2_ fold change (x-axis) and log_10_ P-value (y-axis) from comparing gene expression of *JAK2*V617F *IDH1*R132S (higher expression marked in darkblue) vs *JAK2*V617F *IDH1* wildtype (WT) (higher expression marked in darkred) cells of a patient with chronic phase MPN (SA2CP). Colored are significant different genes (p<0.05) associated with msigdb hallmark signatures and signatures of interest: TNFA signaling via NFKB (TNF; green), Hypoxia (purple), epithelial to mesenchymal transition (EMT; blue), heme metabolism (Heme; red), E2F targets (yellow) and Leukemia stem cell and progenitor-primed (LSPC-primed; black). Genes up regulated in IDH-mutant de novo AML are indicated as *. (C) Heatmap of gene set normalized enrichment scores (NES) from the comparison of the average fold changes *JAK2*V617F *IDH1*R132S vs *JAK2*V617F *IDH1*WT cells of a patient with chronic phase MPN (SA2CP) in key transcriptional compartments. Darkblue indicates higher NES in *JAK2*V617F *IDH1*R132S while dark red indicates higher NES in *JAK2*V617F WT. Columns indicate key transcriptional compartments in CD34+ namely hematopoietic stem cells and lympho-myeloid multipotent progenitor (HSC/LMPP), megakaryocyte/erythroid-primed (Mega/Ery-Primed) and granulocytic/monocytic-primed (Gran/Mono-Primed). False discovery rate (FDR) significance levels are indicated as follows *<0.05; **<0.01; ***<0.001; ****<0.0001. (D) Volcanoplot displaying the log_2_ fold change (x-axis) and log_10_ FDR (y-axis) from comparing gene expression of a patients *IDH1*R132S mutant cells in chronic phase MPN (SA2CP, higher expression marked in darkgreen) vs leukemia transformation (SA2LT; higher expression marked in yellow). Colored and marked are top 70 significant different genes (FDR<0.05) as described in (B). (E) Heatmap of gene set NES from the comparison of the average fold changes of *IDH1*R132S cells in a patient at chronic phase MPN (SA2CP) and after leukemia transformation (SA2LT) in key transcriptional compartments. Darkgreen indicates higher NES in SA2CP while yellow indicates higher NES in SA2LT. Columns indicate key transcriptional compartments in CD34+ namely HSC/LMPP, Mega/Ery-Primed and Gran/Mono-Primed. FDR-significance levels are indicated as follows *<0.05; **<0.01; ***<0.001; ****<0.0001.

Interestingly, even within chronic phase disease, the gene expression changes in *IDH1*-mutant cells showed similarities to gene expression changes in *IDH*-mutant de novo AML^30^, also revealing characteristics such as leukemia stem cell priming and quiescence^27,31^. This occurred alongside the suppression of genes involved in heme metabolism, E2F-targets, DNA metabolic processes, and replication pathways. However, when comparing matched *JAK2*V617F and *IDH1*R132S co-mutated cells between chronic phase disease and post-MPN AML (Figure 6A), we identified more pronounced alterations across all HSPC compartments (Figure 6D-E), including that TNFα signaling and de novo *IDH*-mutant core pathways were upregulated in the leukemic phase, while E2F-targets and DNA replication processes were suppressed. Of note, we observed significant upregulation of TP53 pathway in the Mega/Ery-Primed compartment and other malignancy associated gene expression^20,29,32,33^.

Through linking mutations to transcriptomic profiles via LOTR-Seq, we provide novel insights into the pathways that contribute to *IDH1*-mediated progression to post-MPN AML, highlighting the striking similarities with de novo *IDH*-mutant AML. These findings argue strongly for further investigations into the therapeutic potential of IDH inhibitors during the chronic phase and post-MPN AML settings^34,35^.

### Hyperactivation of JAK-STAT signaling is maintained in *JAK2*V617F-negative post-MPN AML

It has been observed that in 16-50% of post-MPN AML cases, the MPN driver is absent at the time of leukemia transformation, even when the original disease contained a mutation in a canonical MPN driver gene ^36,37,6^. The origin of this MPN driver-negative post-MPN AML is incompletely understood. We reasoned that this category of post-MPN AML must have a different mechanism of transformation compared to the classical linear clonal evolution model.

To examine this further, we used LOTR-Seq to analyze paired, sequential samples from a patient who presented with *JAK2*V617F CP-MPN and *JAK2*V617F-negative post-MPN AML (Figure 7A-H, Supplemental Figure 6A-D). Here, the CP-MPN contained a dominant clone with *TP53*Y236C mutations in addition to a *JAK2*L393V mutation, previously described to result in gain of function^38^ (Figure 6C, Supplemental Figure 7B,D). Within the *TP53*Y236C, *JAK2*L393V population, there was further heterogeneity with two identifiable sub-clonal populations containing either *TP53*R282W (Figure 7D) or *U2AF1*S34Y and *JAK2*V617F (Supplemental Figure 6C; Figure 7B). With evolution to AML, the *TP53*Y236C, *JAK2*L393V *TP53*R282W containing cells demonstrated selective clonal advantage and are dominant in the post-MPN AML stage (Figure 7H). Both *TP53* mutations identified are located within the DNA-binding domain (Supplemental Figure 7A), and thus are predicted to confer a loss-of-function phenotype. Importantly, we were also able to demonstrate that the *TP53* mutations occurred on independent alleles within the same cell and exhibited random monoallelic expression (i.e., somatically acquired in trans; Supplemental Figure 7B-C). These findings identify leukemic transformation that arose from a genetically complex, high-risk ancestral clone and demonstrate parallel clonal evolution resulting in *JAK2*V617F-negative AML (Figure 5I; Supplemental Figure 5E). Here, the hyperactivation of JAK-STAT signaling evident in the chronic phase sample was maintained post leukemic transformation, despite the loss of the *JAK2*V617F clone (Supplemental Figure 7E-F). Of note, maintained hyperactivation of JAK-STAT signaling post leukemic transformation was also observed in the case of linear evolution, patient SA2 (Supplemental Figure 7E-F).

**Figure 7.**
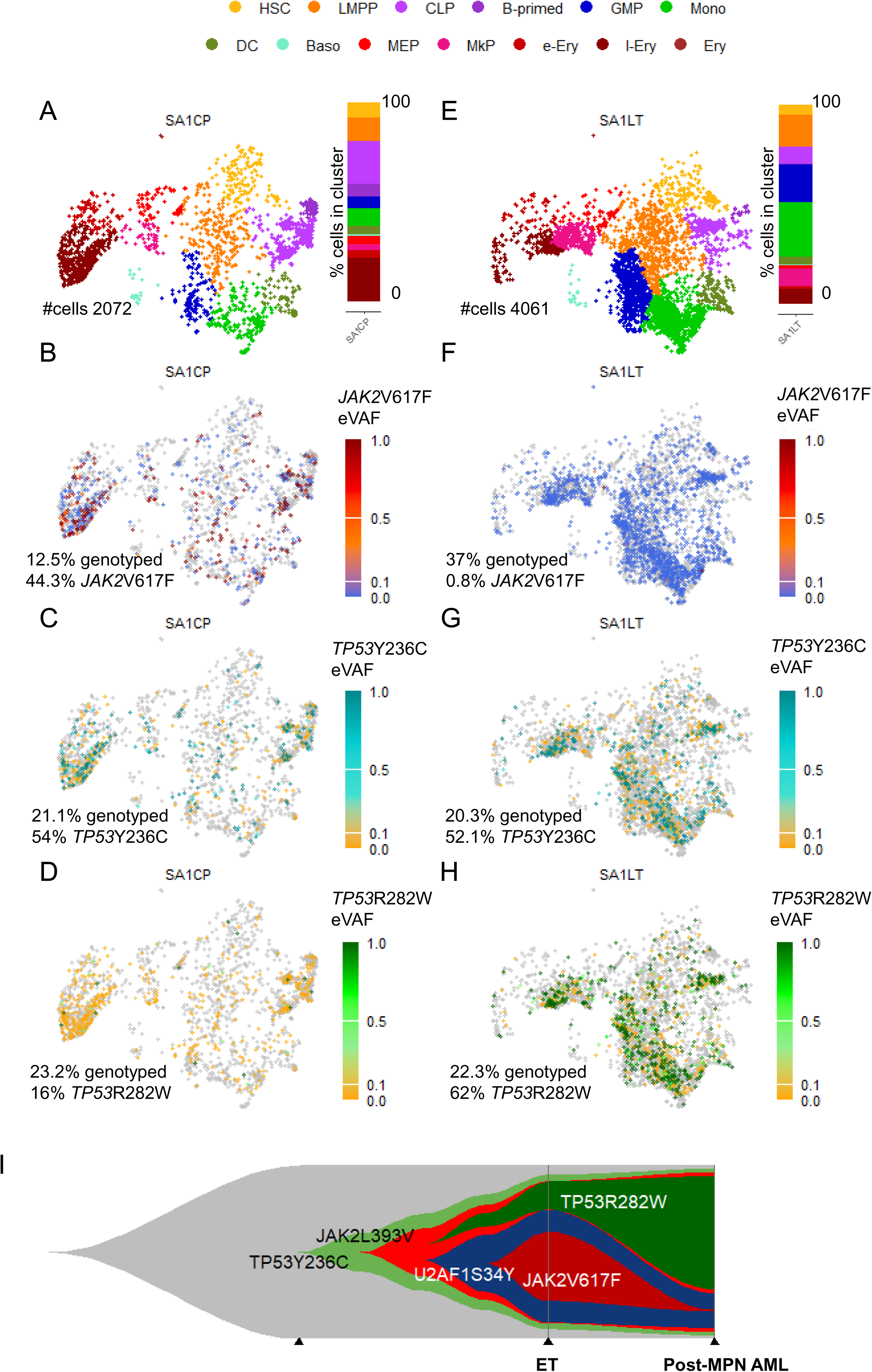
**Leukemic transformation from chronic phase MPN showing parallel clonal evolution of a *JAK2*V617F*-negative TP53-multi-hit* clone.** UMAPs of the same patient at diagnosis of (A-D) chronic phase MPN (SA1CP) and (E-H) post-MPN AML (SA1LT), showing (A,E) transcriptionally defined clusters and bargraphs with the percentage of cells in the clusters and (B-D,F-H) expressed variant allele frequencies (eVAF) of mutations (B,F) *JAK2*V617F, (C,G) *TP53*Y236C and (D,H) *TP53*R282W. Grey colored dots indicate non-genotyped cells. (I) Fish plot illustrating the parallel clonal evolution of the *JAK2*V617F and *TP53*R282W mutant clones.

In summary, using LOTR-Seq to combine single cell transcriptional analysis and panel-enriched genotyping of transcripts from loci commonly associated with myeloid malignancies, we have been able to demonstrate that the transformation of CP-MPN to post-MPN AML is associated with the acquisition of recurrent secondary mutations that drive loss of transcriptional heterogeneity and the immortalization of a genotypically defined, transformed progenitor cell population.

## Discussion

Mutational complexity in MPN is common, with more than 40% of CP-MPN patients harboring a mutation relevant to myeloid malignancies in addition to an MPN driver. Importantly, the acquisition of additional mutations in CP-MPN is associated with reduced event-free survival. Consistent with this, AML arising from antecedent MPN is associated with 2 or more genetic mutations in the vast majority of patients^6^, strongly supporting the hypothesis that the acquisition of these additional mutations drives leukemic transformation of CP-MPN. However, our capacity to determine the functional consequences of mutational complexity in MPN stem cells has been limited by our ability to multiplex single cell phenotyping technology with mutational status. Specifically, single cell RNA-sequencing approaches have provided a high-resolution map of the substantial amount of transcriptional heterogeneity within the HSPC compartment in the BM of patients with CP-MPN but have had limited capacity to associate these differences with an individual cell’s genotype across the spectrum of loci recurrently mutated in post-MPN AML^9,10,22^.

Here, we describe a novel pipeline, LOTR-Seq, combining 10x single cell RNA-sequencing of CP-MPN and post-MPN AML, together with broad, panel-based enrichment and long read genotyping of the vast majority of genetic loci frequently mutated in MPN. We demonstrate that cells expressing the MPN-driver *JAK2*V617F can exhibit transcriptional lineage priming diversity similar to that of their healthy counterparts, but are heavily biased towards a megakaryocyte/erythroid program. Strikingly, this transcriptional diversity is lost upon leukemic transformation, with the HSPC compartment of post-MPN AML exhibiting a homogenous transcriptional profile that appears to be dictated by the identity of the co-occurring mutations.

In the case of the *JAK2*V617F-positive, *TP53*-mutant AML, compound mutant cells were found almost exclusively within the megakaryocyte/erythroid-primed clusters, corroborating recent findings in additional human samples and mouse models with this same genetic combination. Notably, however, the *JAK2*V617F and *IDH*-mutated AMLs exhibited transcriptional features of more primitive HSCs, suggesting that post-MPN is transcriptionally diverse. Testing this hypothesis using a bulk RNA sequencing dataset with a larger number of patients, we demonstrated that post-MPN AML comprises a group of leukemias with three transcriptionally distinct profiles. Importantly, the identity of these transcriptional profiles both diverges and overlaps with what is observed in de novo AML.

Patients with post-MPN AML are excluded from AML clinical trials for targeted agents, attributed to the comparatively dismal outcomes in post-MPN AML. The finding that *IDH*-mutated post-MPN AMLs exhibit transcriptional similarity with the corresponding de novo AMLs should warrant reconsideration of this approach. This is particularly relevant for IDH inhibitors, as the transcriptional consequences of mutant *IDH1* appear to be amplified as either a cause or consequence of leukemic transformation to post-MPN AML.

The progression of CP-MPN to post-MPN AML is rare, difficult to predict, and can occur over decades. Consequently, clinical studies evaluating new MPN therapies are often underpowered to use leukemic transformation as an endpoint. Given the association between mutational complexity and reduced event-free survival, monitoring the emergence of compound mutational clones could prove valuable and should be investigated as surrogate endpoint for CP-MPN disease progression. LOTR-Seq improves upon conventional amplicon-based genomic mutational profiling here by demonstrating conclusively whether TP53 mutations are monoallelic or biallelic, which is highly predictive of prognosis in CP-MPN. LOTR-Seq has also enabled the identification of additional mutations in the genes of interest that are not routinely assessed for in MPN. The identification of *JAK2*L393V has obvious implications for our understanding of, and the evolution of, MPN driver-negative post-MPN AML and CP-MPN.

Furthermore, the additional dimension of transcriptional profiling to the clonal analysis of MPN disease progression allows for both the monitoring of the emergence of high-risk mutations and the assessment of their functional consequence. This information could be used to both inform treatment decisions and allow updated prognostication. It also enables the identification of unique features of high-risk clones that can hopefully be leveraged into new treatments that can prevent the development of post-MPN AML.

In aggregate, the development of LOTR-Seq has both advanced our understanding of the functional consequences of mutational complexity associated with CP-MPN disease progression and provides a new tool to be exploited in CP-MPN clinical trial design to progress the field towards surrogate endpoints of disease progression and ultimately improve long-term outcomes for these patients.

## Acknowledgement

We are grateful for the assistance of the QIMR Berghofer facilities, including flow cytometry and sequencing, and for the helpful comments from members of the Lane Lab, particularly Stacey Anderson. We thank Rachel Thjissen for the fruitful discussion on ONT barcode recovery, primer sequences, and ONT read extension protocols. We gratefully acknowledge the support of Neil Herron and the Herron Family Trust, and the Gordon and Jessie Gilmour Family Trust as dedicated supporters of leukemia research in Queensland. We gratefully acknowledge the contribution of the MPN Alliance Australia for the valuable patient perspective they provided for this study. SWL was funded by an NHMRC Investigator Grant (1195987 2021-2025) and by a philanthropic donation from the MPN Alliance of Australia. ID was supported by a PROMOS, DAAD stipend. JS was funded by a Cancer Council Queensland fellowship (2025829). JG was supported by an HSANZ/Leukaemia Foundation PhD scholarship. This research was supported by the Australian Cancer Research Foundation Centre for Optimised Cancer Therapy.

## Authorship

SWL, MJB, JS conceptualized, designed, and supervised wet lab experiments and bioinformatics analysis. JG, LC, RH, RZ performed wet lab experiments. JS, JG, ID, MB performed bioinformatic analysis. CM, DR, SWL coordinated primary patient sample collection. GA, HC, WG coordinated the collection, consent, and storage of patient sample information. VYL helped with gene panel design. AP performed genomic DNA NGS analysis of primary patient samples. JG, JS, MJB, SWL wrote the manuscript. All authors contributed to and edited the manuscript.

## Data Availability Statement

The 10x sequencing raw and processed data generated in this study is deposited in ArrayExpress with accession E-MTAB-15981. Processed data will be made available via figshare/zendo. Code is available via GitHub https://github.com/JStrau/LOTRSeq. ONT, raw data, due to its size, will be made available upon request.

---

**Supplemental Figure 1.**
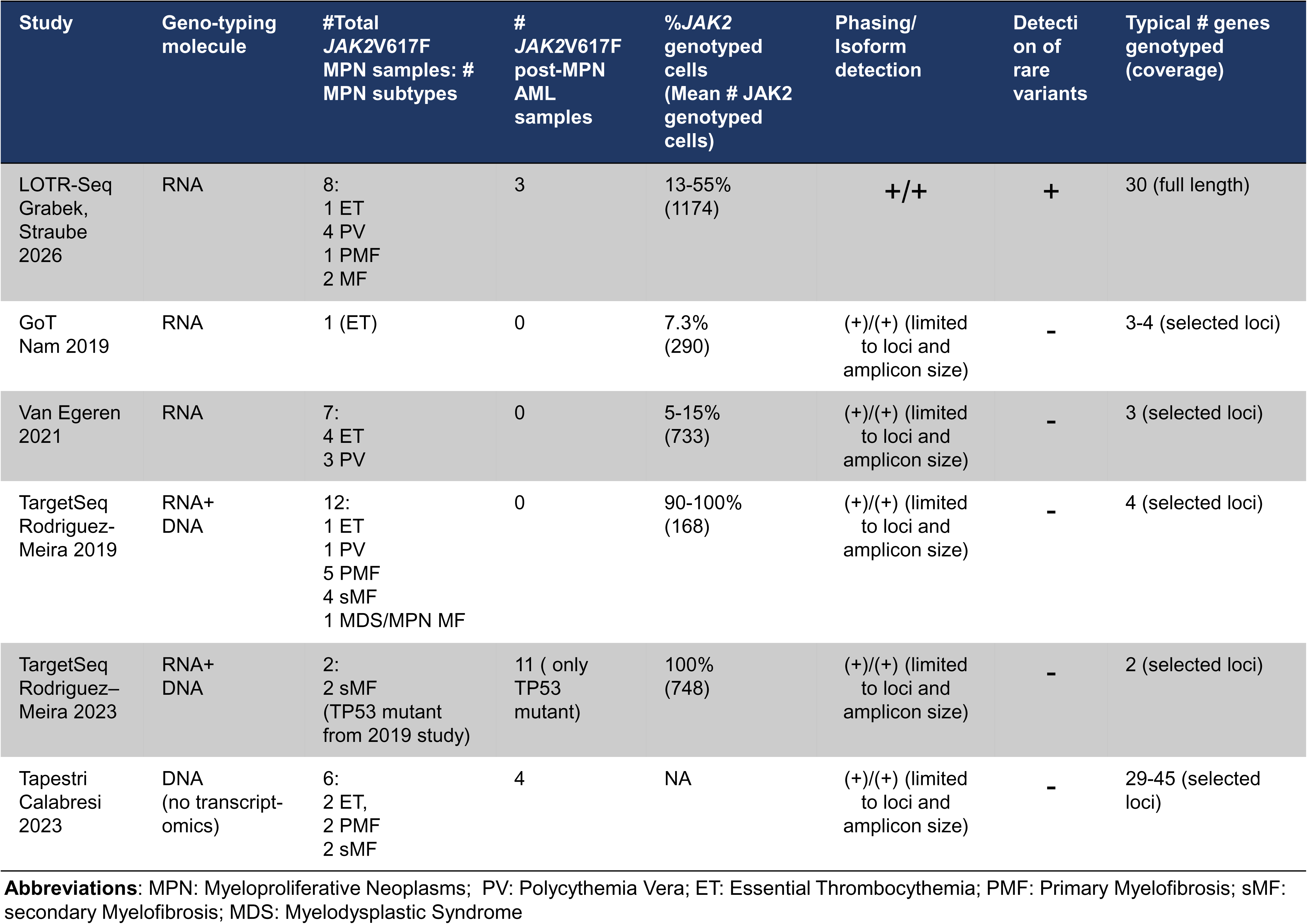

**Supplemental Figure 2.**
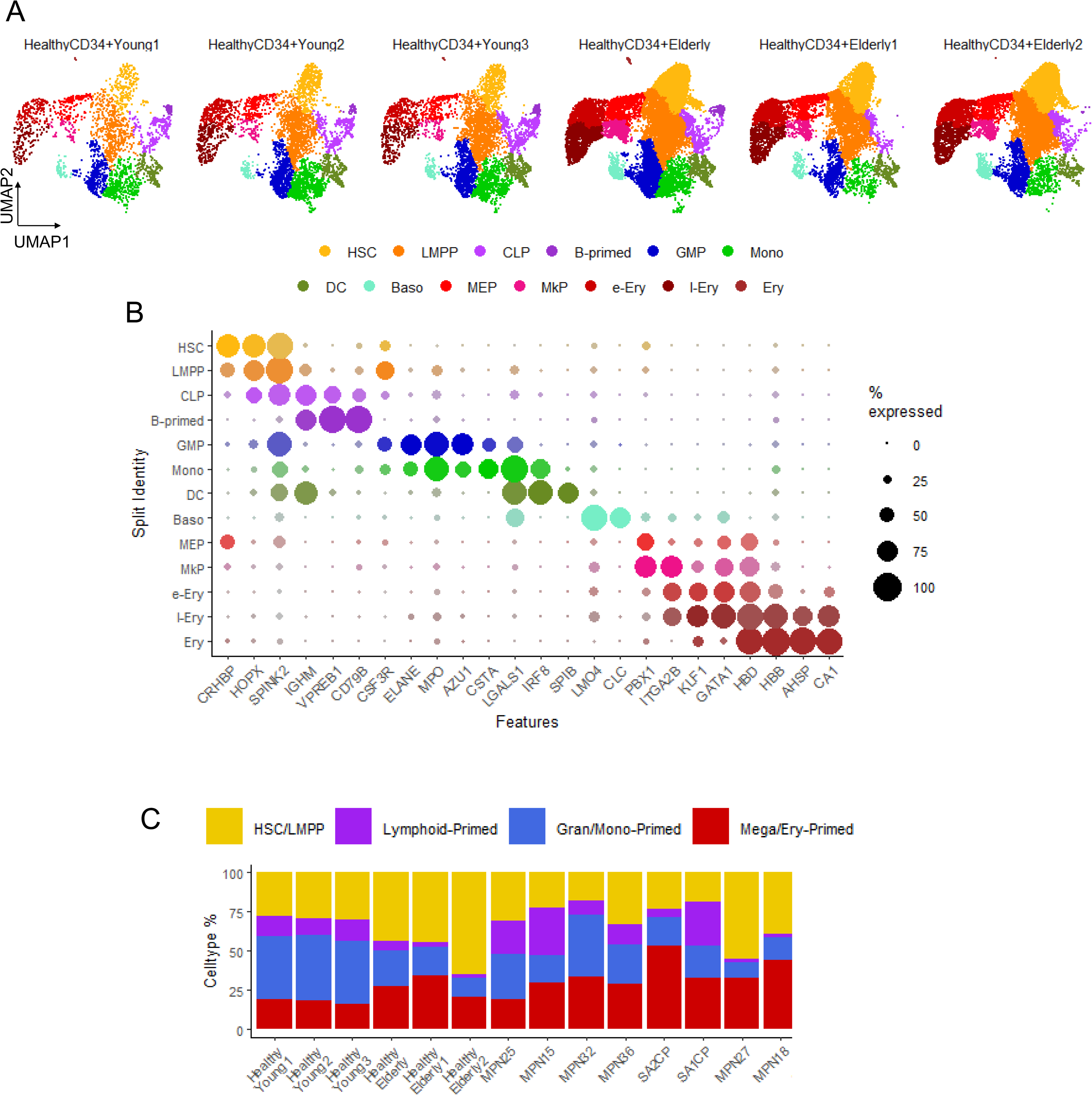

**Supplemental Figure 3.**
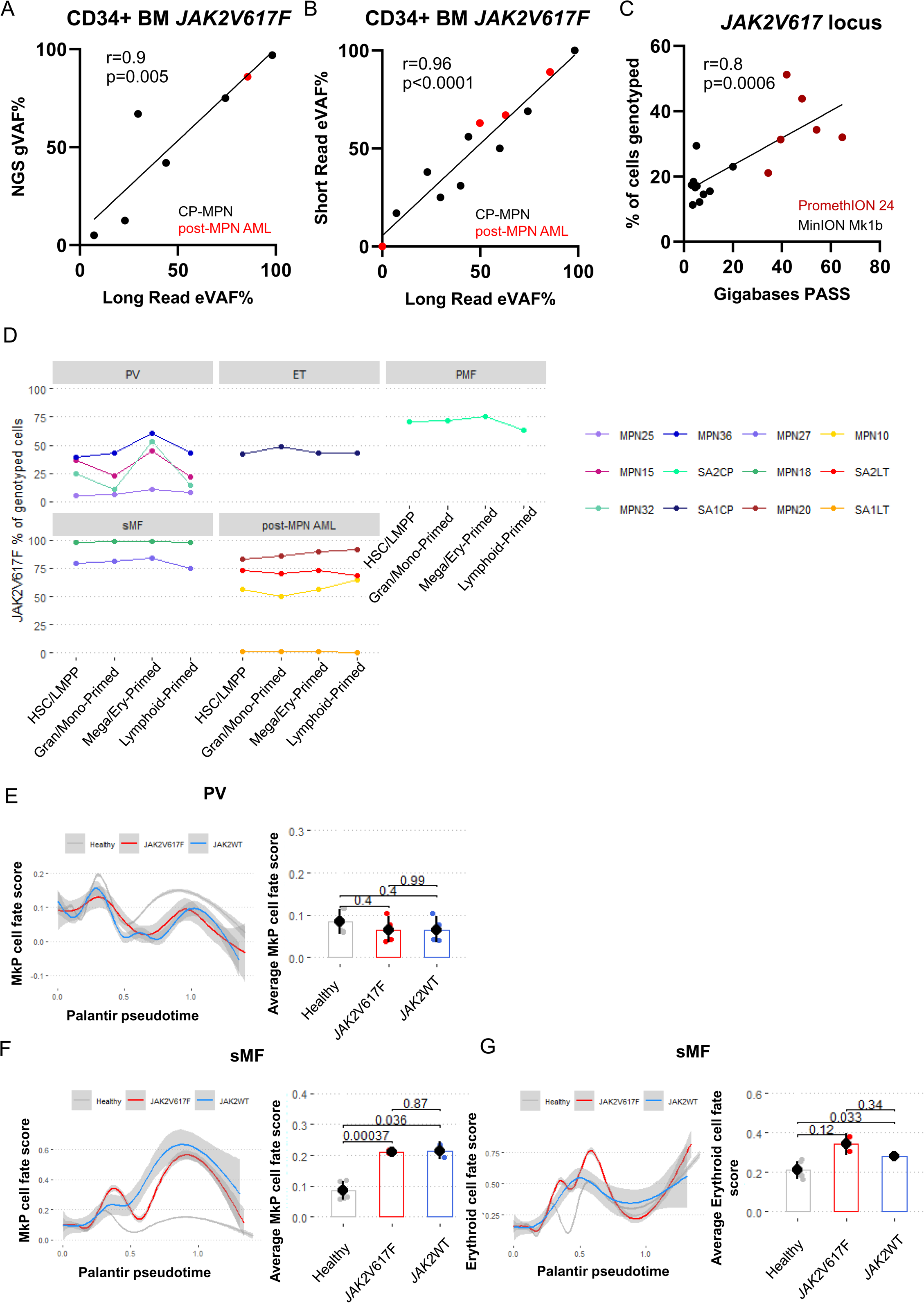

**Supplemental Figure 4.**
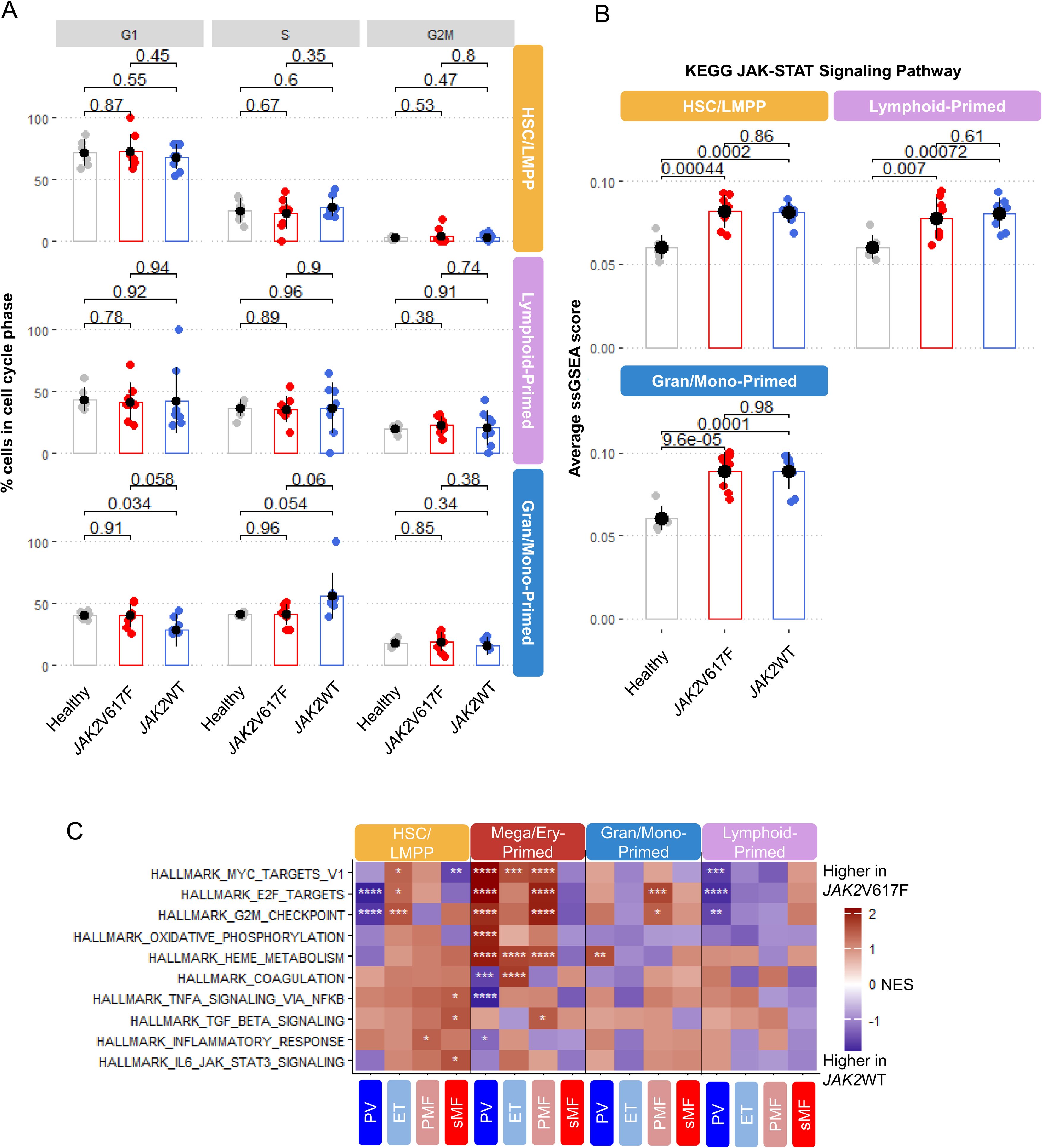

**Supplemental Figure 5.**
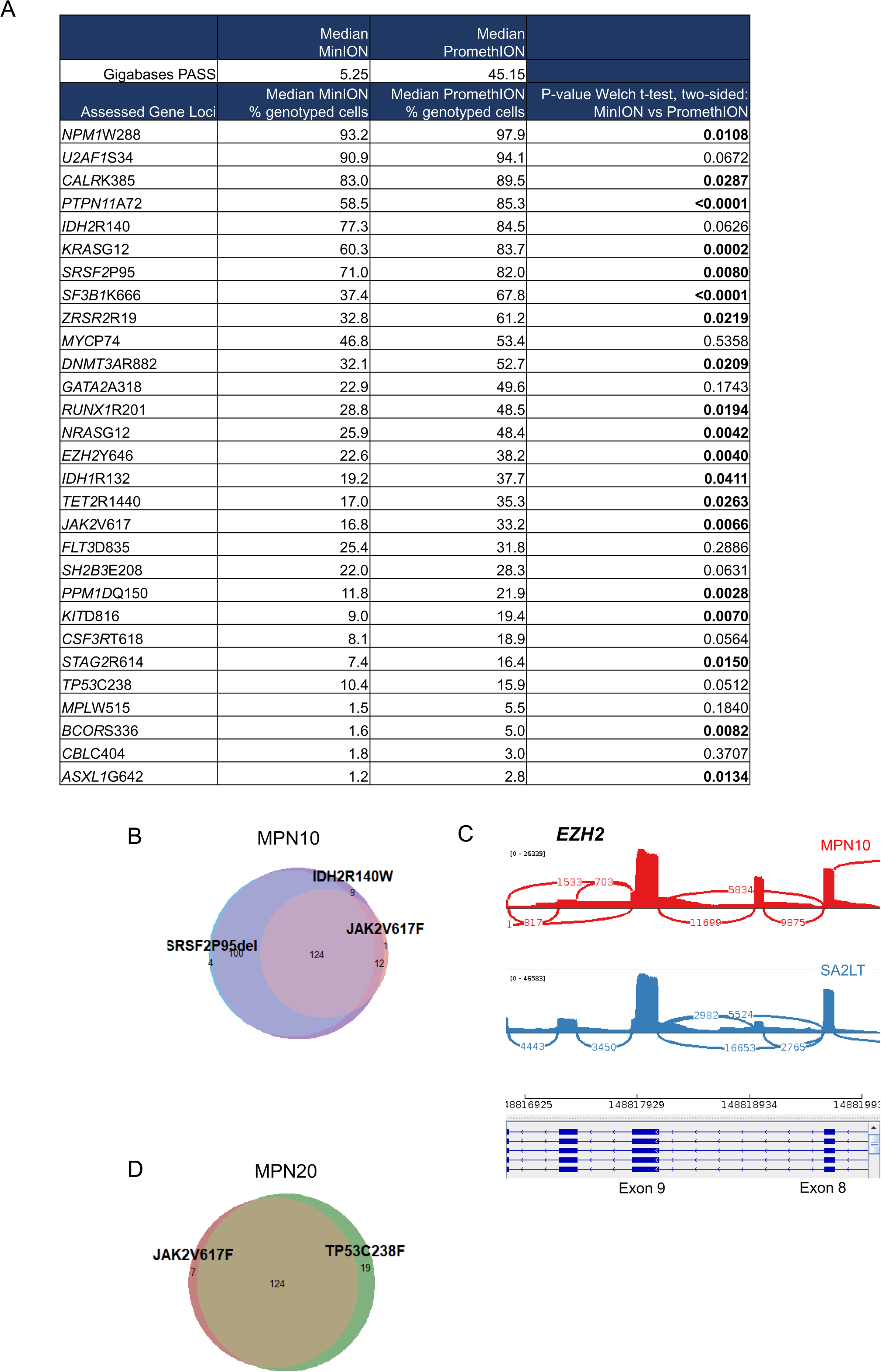

**Supplemental Figure 6.**
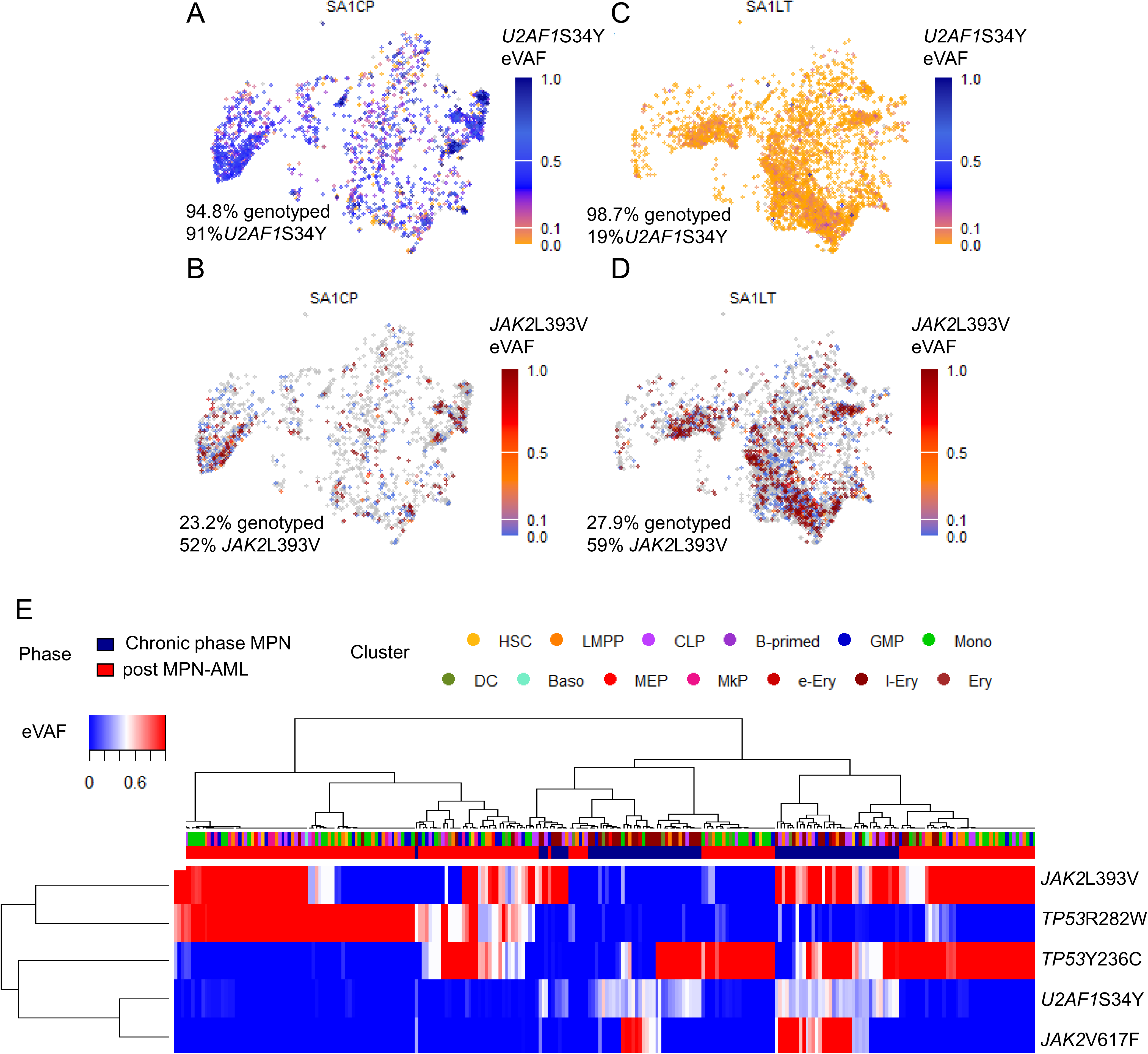

**Supplemental Figure 7.**
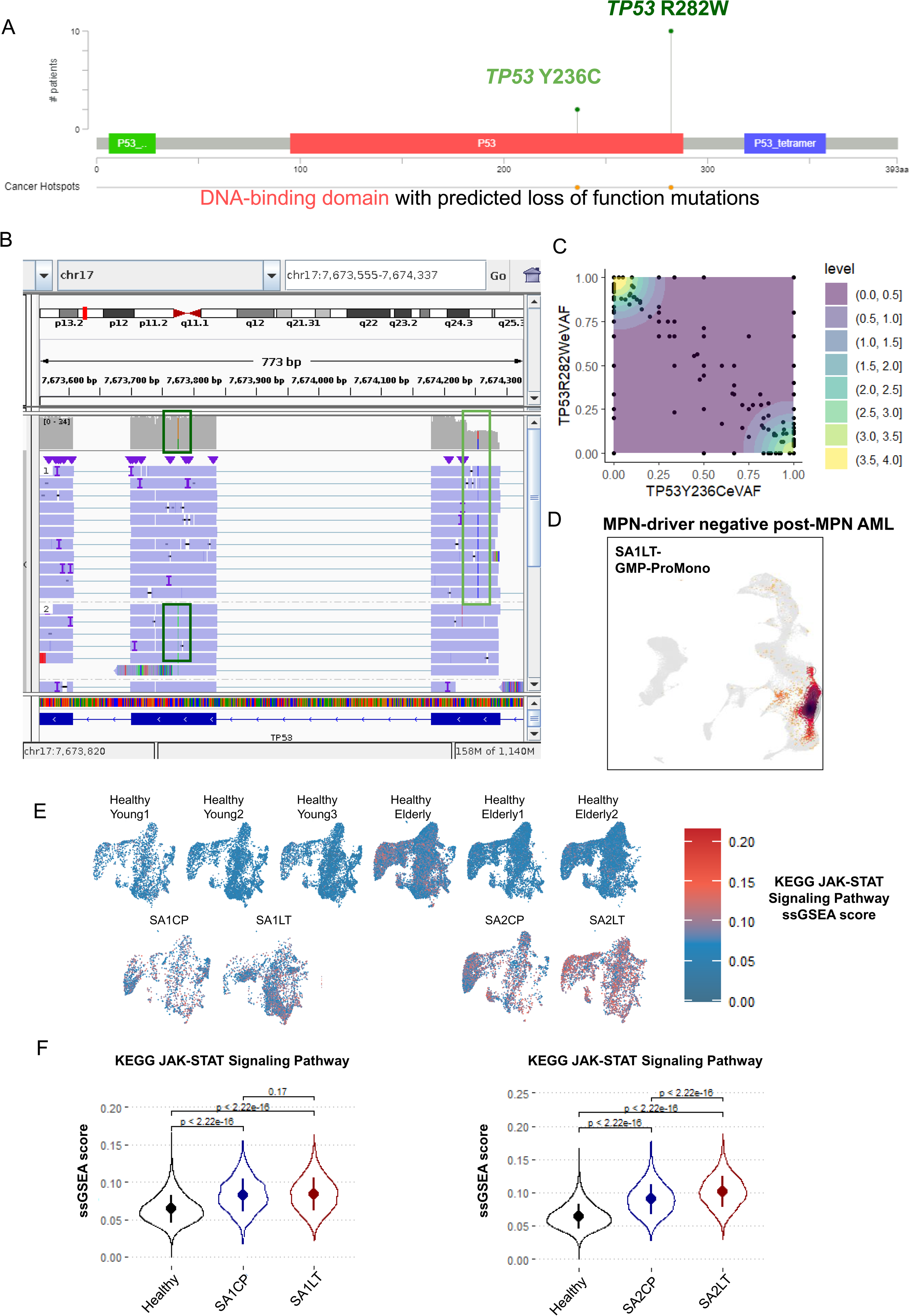

