## Supplemental Material for "Single cell long read genotyping of transcripts reveals discrete mechanisms of clonal evolution in post-MPN AML"

### SUPPLEMENTARY MATERIALS AND METHODS

#### Fluorescence Activated Cell Sorting (FACS) Flow Cytometry

Primary human samples were suspended in staining buffer (PBS with 2% FCS and Pen-Strep). Fluorochromes CD3-PE/Cy7, CD34-PE, CD14-FITC and CD15-APC (supplemental Table 2) were added to the sample at 1µL per $1x10^6$ cells, in order to separate T-cells, hematopoietic stem and progenitor cells, mature monocytes and mature myeloid cells, respectively.

Excess antibody was removed with washing and the sample was re-suspended in sorting buffer (PBS with 2% FCS, PS and 1:20,000 Sytox Blue). Cell Sorting was performed on a BD FACSAria Cell Sorter (BD Biosciences). Following cell sorting, a cell suspension of 1000-1200 cells/µL was used for 10x Chromium Single Cell Encapsulation.

#### Pre and Post-capture PCR Amplification of cDNA

PCR amplification was performed to generate sufficient starting template for target enrichment (based on manufacturer recommendation of minimum of 1000ng). The PCR reaction mix contained 30µL of 10x barcoded cDNA sample, 10µL of partial TSO (5µM), 10µL of R1 primer (5µM) (supplemental Table 3) and 50µL of KAPA HotStart HiFi ReadyMix (Roche, Ref 07958927001).

Pre-capture PCR was performed with the following incubation times of 98°C for 3 mins then the PCR cycle commences at 98°C for 20 secs followed by 63°C for 30 secs then 72°C for 4.5 mins. The cycle number varied based on starting amount with between 12-24 cycles required. The final extension step was 72°C for 3:00 mins.

The sample underwent AMPure Bead clean up at 1X with 80% Ethanol. The cDNA was eluted into PCR grade water and quantified by Qubit (Invitrogen, ThermoFisher Scientific).

The post capture PCR Master Mix contains 25µL of KAPA HiFi HotStart ReadyMix (2X) with 2.5 µL of 5µM R1 primer and 2.5µL of 5µM partial TSO primer (supplemental Table 3). The 30µL of master mix was added to the 20µL of bead bound cDNA from the post capture bead wash.

**Single cell short read sequencing processing**

In brief, cells were retained if their mitochondrial RNA content was below 15% and they had more than 200 expressed genes. Each sample was normalized, and the top 2000 variable features identified. For sample integration, the top 2000 integration features were detected (`SelectIntegrationFeatures()`), followed by identification of the top 2000 integration anchors (`FindIntegrationAnchors()`). The anchors were then used as input for the `IntegratedData function` with default parameters. Integrated data were cell cycle scored (‘CellCycleScoring()’). Uniform Manifold Approximation and Projection (UMAP) dimensional reduction was performed. Clustering was performed using the shared nearest neighbor graph construction (`FindNeighbors`) on the first two UMAP components, followed by optimization the modularity function (`FindClusters`) with a resolution of 0.1.

**ONT Base calling and chimeric read separation**

Raw fast5 signals were converted using ONT’s pod5 Python package (v0.1.21). Basecalling was performed with ONT’s dorado basecaller (v0.7.1) with `--no-trim` option using transformer models specific for the sample Ligation Kit and Platform detailed in supplemental Table 4. Chimeric read trimming was perform using porechop (v0.2.4; https://github.com/rrwick/Porechop) with custom adapter settings added to adapter.py code ` Adapter('PCR adapters 10x', start_sequence=('PCR_10x_start', 'CTACACGACGCTCTTCCGATCT'), end_sequence= ('PCR_10x_end', 'AGATCGGAAGAGCGTCGTGTAG')) and settings `--extra_middle_trim_bad_side 0 --min_split_read_size 200 --extra_middle_trim_good_side -25 --extra_end_trim -25`.

**Differential gene expression and gene set enrichment analysis**

Differential gene expression was performed with edgeR (v4.6.3). Single cell gene set enrichment analysis (ssGSEA) was performed on normalized counts with GSVA (v1.52.3) R package. Gene set enrichment analysis on edgeR derived log fold changes was performed using fgsea18 (v1.34.2) with default parameters. Gene sets were downloaded MSigDB (v2025.1). IDH mutant AML core gene set was generated using Sirenko et al. 2025 extracted TableS3 data by intersecting differential up or down regulated genes (p_val_adj<0.05) across cell types (supplemental Table 6).

**Mutation and INDEL calling**

For single nucleotide variant (SNV) calling, we used bcftools (v1.19) mpileup function on processed bams to count bases at gene loci of interest (supplemental Tables 7-8). Insertion/Deletion (INDEL) calling was performed only over known mutation regions using samtools (v1.20) mpileup with `-d 0` option and VarScan2 (v2.4.2)[1] mpileup2indel with options` --min-coverage 0 --min-reads2 0 --min-avg-qual 0 --output-vcf 1 --variants 1`.

### SUPPLEMENTARY FIGURE LEGENDS

**Supplementary Figure 1.** Table comparing published single cell genotyping methods applied to MPN samples.

**Supplementary Figure 2.** Lineage-defining gene expression markers in CD34+ HSPCs. (A) UMAPs of n=6 healthy CD34+ bone marrow single cells derived from short read sequencing whole transcriptomics (B) Dotpot of average percentage of lineage defining markers expressing cells in healthy samples (n=6, x-axis) within the annotated clusters (y-axis). (C)  Bargraph of the relative cell cluster percentage (y-axis) of each analyzed sample (x-axis; n=6 healthy; n=8 chronic phase MPN).

**Supplementary Figure 3.** Scatter plots with linear regression lines of (A-C) Nanopore long read expressed variant allele frequency (eVAF; x-axis) compared to (A) the next-generation sequencing derived (NGS) genomic VAF (gVAF; y-axis) and (B) pseudobulked eVAF derived from short-read sequencing (y-axis) and, (C) Nanopore sequenced gigabases PASS (x-axis) and the percentage (%) of genotyped cells (y-axis). Statistics displayed are the Pearson's product-moment correlation coefficient r and test for association derived p-value. (D) Scatter plot showing the *JAK2*V617F percentage (%) of genotyped cells (y-axis) per cellular compartment (x-axis) for each patient separated by MPN subtype or post-MPN AML. (E-G) Smoothed regression lines of the Palantir predicted (E-F) megakaryocyte progenitor (MkP) or (G) erythroid cell fate score (y-axis) in dependency of the pseudotime (x-axis) and bargraph of the average

Palantir scores for n=5 healthy, n=4 Polycythemia vera (PV; E) or n=2 secondary myelofibrosis (sMF; F,G) of *JAK2*V617F and JAK2-wild type (JAK2WT) cells. In the bargraphs black dot and line represent the group mean and standard deviation, respectively. P-values derived from pairwise, two-sided Welch t-tests.

**Supplementary Figure 4.** (A) Bargraph of the percentage of cell cycle phase (G1, S, G2M) for each compartment comparing n=6 healthy to n=8 MPN *JAK2*V617F and *JAK2*-wild type (*JAK2*WT). Black dot and line represent the group mean and standard deviation, respectively. P-values derived from pairwise, two-sided Welch t-tests. (B) Bargraph of the KEGG JAK-STAT signaling pathway average single cell gene set enrichment (ssGSEA) score comparing n=6 healthy to n=8 MPN *JAK2*V617F and *JAK2*-wild type (*JAK2*WT). Black dot and line represent the group mean and standard deviation, respectively. P-values derived from pairwise, two-sided Welch t-tests. (C) Heatmap of the gene set enrichment analysis of selected mSigDB Hallmark pathways comparing *JAK2*V617F mutant cells to JAK2-wild type (WT) cells for patient with Polycythemia Vera (PV; n=4), Essential Thrombocythemia (ET;n=1), primary myelofibrosis (PMF,n=1), and secondary myelofibrosis (sMF,n=2), separated by lineage primed clusters. The normalized enrichment score (NES) is colored red if the hallmark is enriched in JAK2V617F cells and blue if it is enriched in JAK2WT. False Discovery Rate (FDR) significance levels are represented as follows FDR **** <0.0001 ; *** <0.001 ; **<0.01; * <0.05.

**Supplementary Figure 5.** (A) Table of the median percentage (%) of genotyped cells at MPN/AML mutated genes hotspot locations for n=10 MinION sequenced samples and n=6 PromethION sequenced samples. P-values derived from a pairwise, two-sided Welch t-test. (B,D) Euler diagrams depicting the number of mutated cells with genotyping information for all 2-3 loci by LOTR-Seq and their co-occurrence for samples (B) MPN10 and (D) MPN20. (C) Sashimi plot showing *SRSF2*P95R target *EZH2* retaining a poison exon cassette in long read sequencing of *JAK2*V617F *SRSF2*P95R *IDH2*R140W mutant post-MPN patient MPN10 but not *JAK2*V617F *IDH1*R140W mutant post-MPN AML patient SA2LT.

**Supplementary Figure 6.** UMAPs of the same patient at diagnosis of (A-B) chronic phase MPN (SA1CP) and (C-D) post-MPN AML (SA1LT), displaying expressed variant allele frequencies (eVAF) of mutations (A,C) *U2AF*1S34Y and (B,D) *JAK2*L393V. Grey colored dots indicate non-genotyped cells. (E) Hierarchical clustering using ward.D2 method of the Euclidean distance of eVAFs of mutated genes *JAK2*L393V, *TP53*R282W, *TP53*Y236C, *U2AF1*S34Y *and JAK2*V617F of SA1CP and SA1LT.

**Supplementary Figure 7.** (A) Lollipop plot extracted from cBioportal depicting the location of *TP53*R282W and *TP53*Y236C in the DNA binding domain of TP53. (B) IGV screen shot of the mapped ONT long reads from SA1LT of a single cell with *TP53*R282W and *TP53*Y236C mutations on independent reads. (C) Scatter and density plot of the expressed variant allele frequency (eVAF) of *TP53*Y236C (x-axis) and *TP53*R282W (y-axis). (D) SA1LT CD34+ HSPC single cell gene expression projected on Zeng et al. 2025 bone marrow atlas UMAP[2]. (E-F) Single cell geneset enrichment analysis (ssGSEA) score of the msigDB KEGG JAK-STAT Signalling pathway (A) visualised on the UMAPs of healthy individuals and two patients during chronic phase (CP) MPN and leukemia transformation (LT). Patient SA1 presented with two *JAK2* mutations affecting amino acids V617F and L393V in independent clones in chronic phase (SA1CP) but progressed with JAK2V617F-negative but JAK2L393V-positive clone post MPN AML (SA1LT). In addition, we also visualized a patient SA2 that has persistent JAK2V617F mutation during chronic phase MPN (SA2CP) and leukemia transformation (SA2LT). (F) Violin plots with mean and standard deviation comparing ssgGSEA scores of n=6 pooled healthy individuals to patient SA1 during CP and LT or patient SA2 CP and LT. P-values derived from pairwise, two-sided Welch t-tests.

1. Koboldt DC, Zhang Q, Larson DE, et al. VarScan 2: somatic mutation and copy number alteration discovery in cancer by exome sequencing. *Genome Res*. Mar 2012;22(3):568-76. doi:10.1101/gr.129684.111
2. Zeng AGX, Iacobucci I, Shah S, et al. Single-cell Transcriptional Atlas of Human Hematopoiesis Reveals Genetic and Hierarchy-Based Determinants of Aberrant AML Differentiation. *Blood Cancer Discov*. Jul 1 2025;6(4):307-324. doi:10.1158/2643-3230.BCD-24-0342

178
